# Recovering high-resolution information using energy filtering in MicroED

**DOI:** 10.1101/2025.03.26.645403

**Authors:** Max T.B. Clabbers, Tamir Gonen

## Abstract

Inelastic scattering poses a significant challenge in electron crystallography by elevating background noise and broadening Bragg peaks, thereby reducing the overall signal-to-noise ratio. This is particularly detrimental to data quality in structural biology, and the diffraction signal is relatively weak. These effects are aggravated even further by the decay of the diffracted intensities as result of accumulated radiation damage, and rapidly fading high-resolution information can disappear beneath the noise. Loss of high-resolution reflections can partly be mitigated using energy filtering, which removes inelastically scattered electrons and improves data quality and resolution. Here, we systematically compared unfiltered and energy-filtered MicroED data from proteinase K crystals, first collecting an unfiltered dataset followed directly by a second sweep using the same settings but with the energy filter inserted. Our results show that energy filtering consistently reduces noise, sharpenes Bragg peaks, and extends high-resolution information, even though the absorbed dose was doubled for the second pass. Importantly, our results demonstrate that high-resolution information can be recovered by inserting the energy filter slit. Energy-filtered datasets showed improved intensity statistics and better internal consistency, highlighting the effectiveness of energy filtering for improving data quality. These findings underscore its potential to overcome limitations in macromolecular electron crystallography, enabling higher-resolution structures with greater reliability.

---

Inelastic scattering poses a significant challenge in electron crystallography, as inelastic events have a 3-4 times higher probability to occur compared to elastic scattering in hydrated protein crystals (Henderson, 1995; Latychevskaia & Abrahams, 2019). Inelastically scattered electrons lose coherence, causing elevated background levels and a broadening of the Bragg spots. This increase in noise can interfere with detection of the diffraction spots and reduce accuracy of the intensity measurements. The effects are particularly detrimental to data quality from biological samples, which typically have a relatively weaker diffraction signal and experience substantial scattering contributions from the bulk solvent. This is further exacerbated during data collection, as the mean diffracted intensities rapidly fade owing to radiation damage (Hattne *et al*., 2018). As a result of the increasing dose, faint high-resolution reflections are lost when the diffraction signal at high scattering angles drops below the background noise. Loss of high-resolution information, due to dose-sensitivity and increased noise from inelastic scattering, compromises the accuracy of structural models and can obscure detailed features essential for biological interpretation. Therefore, mitigating these effects is crucial for improving data quality and achieving more precise structural insights.

Energy filtering removes inelastically scattered electrons above a defined energy-loss threshold, improving the signal-to-noise ratio and quality of the diffraction measurements in both materials science (Gemmi & Oleynikov, 2013; Yang *et al*., 2022) and structural biology (Gonen *et al*., 2004, 2005; Yonekura *et al*., 2002, 2015, 2019). Recently, we implemented a strategy for microcrystal electron diffraction (MicroED) data collection that combines electron counting (Martynowycz *et al*., 2022) with energy filtering (Clabbers *et al*., 2025a). Using this integrated approach, energy-filtered data from proteinase K crystals showed a significant improvement in overall data quality and high-resolution information extending to 1.09 Å, compared to 1.4 Å resolution for crystals recorded without energy filtering (Clabbers *et al*., 2025a). Further analysis of energy filter slit widths revealed narrower slit settings reduced noise and yielded sharper peaks, improving spot separation and intensity measurements with better internal consistency, ultimately leading to more accurate structural models (Clabbers *et al*., 2025b). However, direct comparisons of intensity and model statistics between filtered and unfiltered data between crystals remain challenging due to variations in crystallinity, isomorphism, and radiation damage between different crystals.

Here, we systematically compare unfiltered and energy-filtered MicroED data from the same crystal samples. Proteinase K crystals were machined using focused ion beam milling to lamellae with an optimal thickness of 300 nm (Martynowycz *et al*., 2021, 2023) and transferred to the transmission electron microscope (TEM) that was aligned with low flux density conditions of ∼0.002 e^-^/Å^2^ for data collection (Clabbers *et al*., 2025a). For each lamella, two consecutive data collection passes were performed. First, an unfiltered dataset was recorded with the energy filter slit retracted, covering a 20.0° continuous rotation sweep with a total fluence of 0.84 e^-^/Å^2^, corresponding to an absorbed dose of ∼3.1 MGy. At this amount of dose evidence of radiation damage should be visible (Garman 2010, Hattne *et al*., 2018). Immediately after, a second pass was recorded using the same protocol, but with the energy filter slit inserted. This approach was repeated for 11 lamellae to ensure sufficiently high completeness and minimize variability for a comparative analysis. To enable a direct comparison, data were processed using XDS and truncated at a cross-correlation between two random half sets (*CC*_1/2_) that was still significant at the 0.1% level in the highest resolution shell (Karplus & Diederichs, 2012).

Consistent with past studies (Clabbers *et al*., 2025a,b), energy filtering visibly improved the MicroED data quality, with reduced background noise, sharper Bragg peaks, and high-resolution information extending out further compared to unfiltered data (**Fig. 1**). Interestingly, the filtered MicroED data consistently showed improved intensity statistics, and in all but two instances higher resolution information, even though the absorbed dose for the second pass was effectively doubled compared to any of the unfiltered datasets (**Fig. 2, S1, and S2**). For example, a first crystal recorded without filtering was truncated at 1.34 Å resolution, with a *CC*_1/2_ of 19.8% and a mean *I/σI* of 0.89 in the highest resolution shell. In contrast, the second energy-filtered pass provided significantly higher information up to 1.06 Å resolution, with a *CC*_1/2_ of 10.2% and a mean *I/σI* of 0.71 (**Fig. 2**). In the two cases where the resolution did not improve, intensity statistics of the filtered data were significantly better in all resolution shells compared to their unfiltered counterparts (**Fig. 2 and S1**). Variations in attainable resolution for different lamellae likely stem from differences in crystallinity, lamella quality, and lattice orientation.

**Figure 1.**
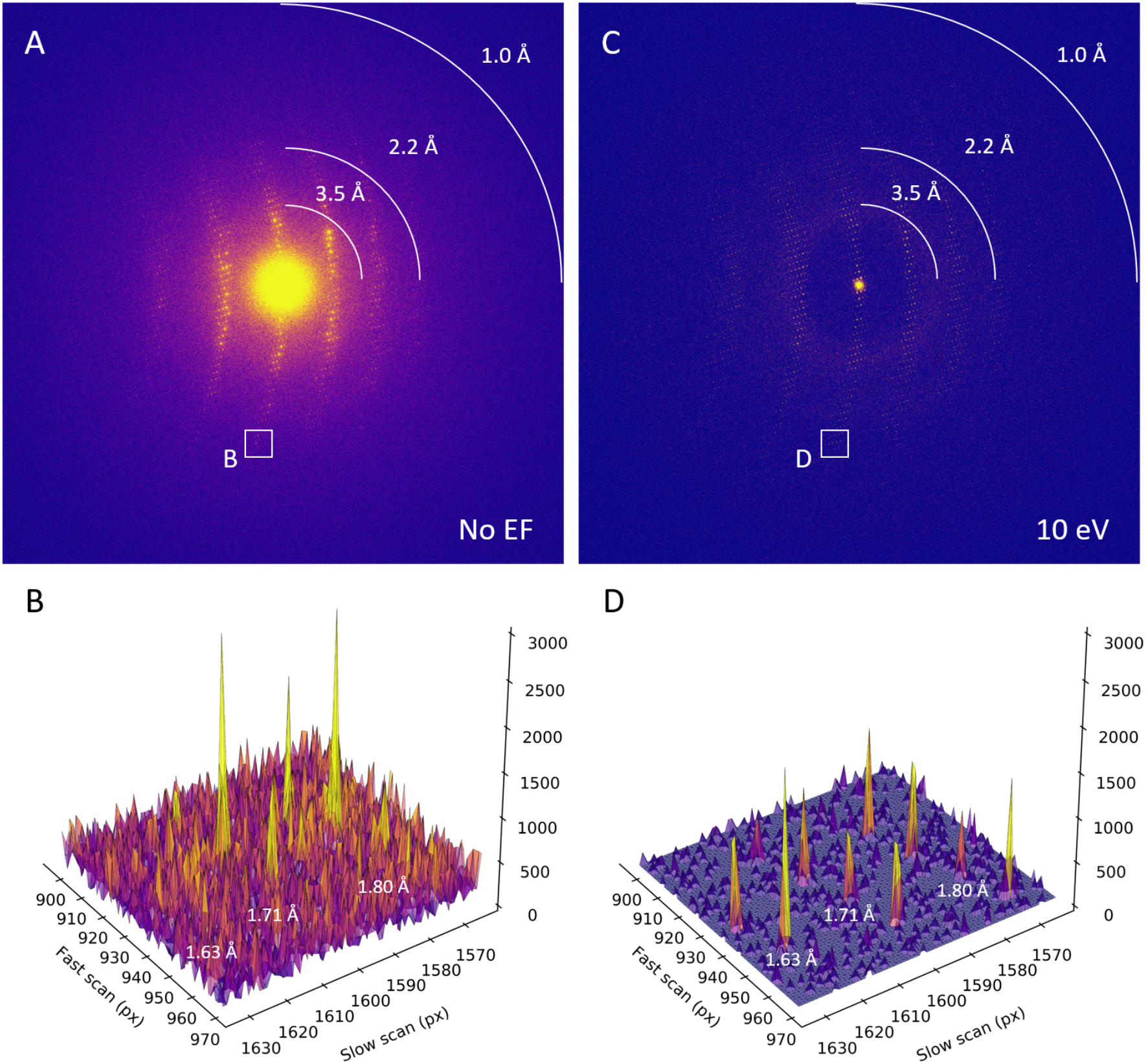
Unfiltered and filtered MicroED diffraction patterns. **A.**Diffraction frame summed over the last 10 s of unfiltered MicroED data collection with an exposure of 0.02 e^-^/Å^2^ for the frame and an accumulated dose of ∼3.1 MGy. Peak profiles for the area highlighted marked with B. are shown in the corresponding subpanel. **B**. Peak profiles from the area highlighted in A. showing rows of diffraction spots at respectively ∼1.6, ∼1.7, and ∼1.8 Å resolution. **C**. Diffraction frame covering the same wedge in reciprocal space as shown in A. for the second energy-filtered MicroED pass, with an exposure of 0.02 e^-^/Å^2^ for the frame and a total accumulated dose of ∼6.2 MGy. Peak profiles for the area highlighted marked with D. are shown in the corresponding subpanel. **D**. Peak profiles for filtered data for the same inset as shown in B. from the area highlighted in D. showing rows of diffraction spots at respectively ∼1.6, ∼1.7, and ∼1.8 Å resolution that have a much stronger separation of the Bragg peaks from the background noise compared to the unfiltered data in B.

**Figure 2.**
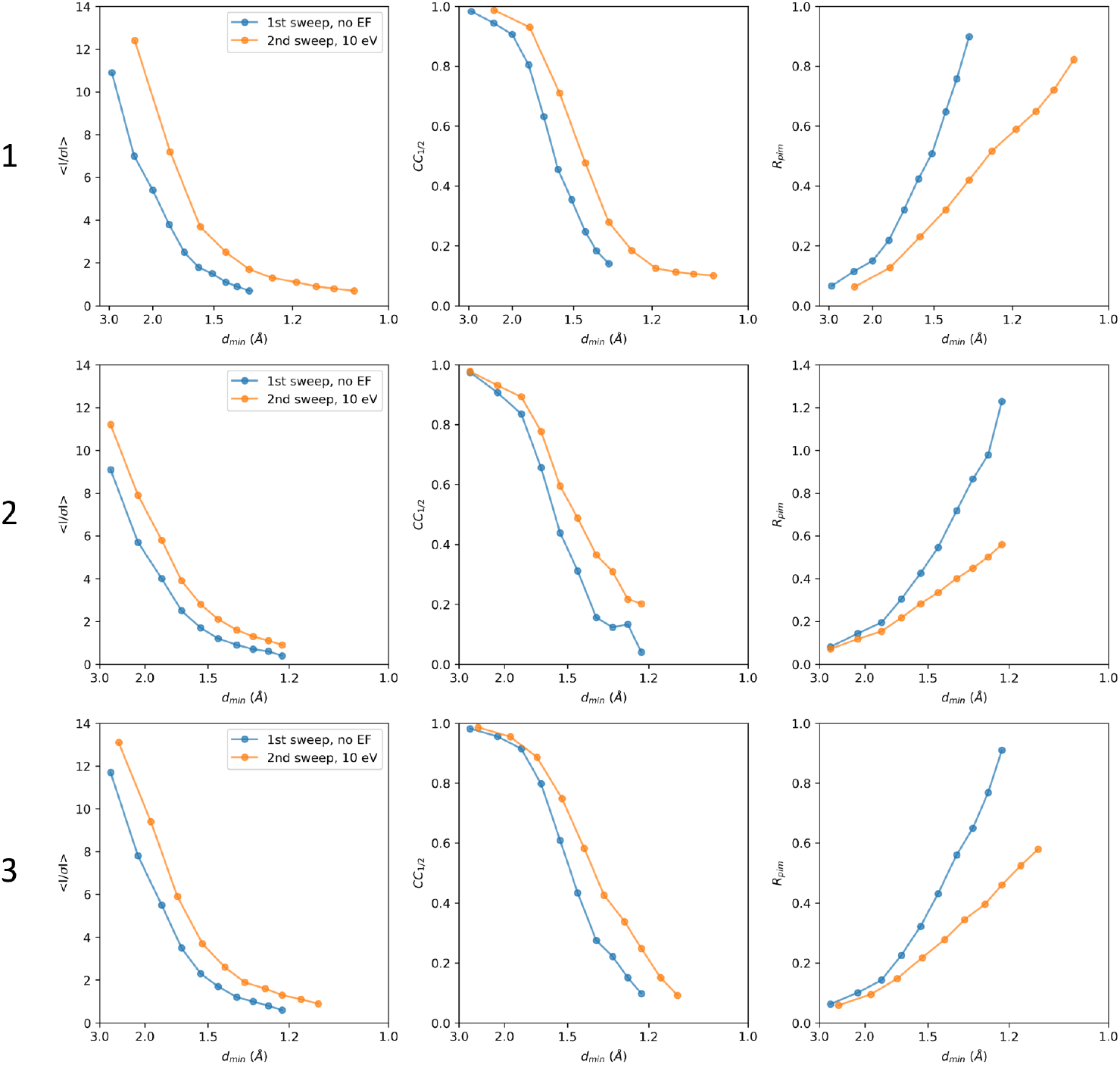
Intensity statistics for three MicroED data collection series. For each series, two datasets were collected from the same lamellae where the first pass did not use energy filtering (blue), whereas the second pass had the 10 eV energy filter slit inserted (orange). For each lamella, the crystallographic quality indicators mean *I/σI, CC*_1/2_, and *R*_pim_ are plotted as function of the resolution. Individual datasets were truncated at a *CC*_1/2_ that was still significant at the 0.1% level in the highest resolution shell.

The merged unfiltered data from the first data collection sweeps showed signification information up to 1.3 Å resolution (**Table 1**). To enable a direct comparison of the intensity and model statistics, data recorded with energy filtering were truncated at the same resolution, even though the crystals diffracted to higher resolution in the second sweep (**Table 1, Fig. 3**). At the same resolution, the energy-filtered data show better statistics such as the mean *I/σI, CC*_1/2_, and *R*-factors, across all resolution shells, despite having absorbed twice the dose compared to the unfiltered data (**Fig. 3**). Notably, the mean *I/σI* in the highest resolution shell was twice as high, and the *CC*_1/2_ was improved overall indicating a better internal consistency between the intensity measurements of the filtered data (**Table 1**). The quality of the resulting models was rather similar, showing a *R*_work_/*R*_free_ of 0.189/0.228 for the filtered data compared to the unfiltered data at 0.181/0.226 (**Table 1**). The fact that the model *R*-factors for the filtered data did not improve, even though the intensity statistics were significantly better, likely is the result of the higher accumulated dose and increased radiation damage compromising structural integrity. Visual inspection of the map did not reveal any major differences in model quality, and *B*-factors were rather similar (**Table 1**).

**Table 1.** Data processing and refinement statistics.

|  | 1<sup>st</sup> pass, no EF | 2<sup>nd</sup> pass, 10 eV EF slit |
| --- | --- | --- |
| **Data collection** |  |  |
| No. of crystals | 11 | 11 |
| Flux density (e<sup>-</sup>/Å<sup>2</sup>/s) | 0.002 | 0.002 |
| Fluence (e<sup>-</sup>/Å<sup>2</sup>) | 0.84 | 0.84 |
| Total fluence (e<sup>-</sup>/Å<sup>2</sup>) | 0.84 | 1.68 |
| Dose (MGy) | 3.08 | 6.16 |
| **Data integration** |  |  |
| Space group | *P*4<sub>3</sub>2<sub>1</sub>2 | *P*4<sub>3</sub>2<sub>1</sub>2 |
| Unit cell dimensions |  |  |
| *a*, *b*, *c* (Å) | 67.62, 67.62, 106.59 | 67.64, 67.64, 106.67 |
| *α*, *β*, *γ* (°) | 90, 90, 90 | 90, 90, 90 |
| Resolution (Å)<sup>†</sup> | 57.10-1.30 (1.33-1.30) | 43.64-1.30 (1.33-1.30) |
| Observed reflections | 1,377,854 (102,897) | 1,348,162 (102,276) |
| Unique reflections | 59,002 (4,294) | 59,010 (4,297) |
| Multiplicity | 23.4 (24.0) | 22.8 (23.8) |
| Completeness (%) | 96.1 (96.3) | 95.9 (96.0) |
| R<sub>merge</sub> | 0.495 (2.463) | 0.302 (1.084) |
| *<I/σI>* | 5.2 (1.0) | 6.7 (1.9) |
| CC<sub>1/2</sub> (%) | 95.9 (15.2) | 98.9 (23.9) |
| **Refinement** |  |  |
| Resolution (Å) | 57.10-1.30 | 43.64-1.30 |
| No. of reflections | 57,511 | 58,442 |
| R<sub>work</sub>/R<sub>free</sub> | 0.181/0.226 | 0.189/0.228 |
| *<B>* factors (Å<sup>2</sup>) |  |  |
| Overall | 12.57 | 12.61 |
| Protein | 9.98 | 10.48 |
| Ligands/ions | 10.68 | 11.87 |
| Waters | 27.62 | 27.35 |
| R.m.s. deviations |  |  |
| Bond lengths (Å) | 0.011 | 0.009 |
| Bond angles (°) | 1.13 | 1.02 |

<sup>†</sup>Values in parenthesis are for the highest resolution shell. Data were truncated at 1.3 Å resolution to allow for a direct comparison of intensity and model statistics.

**Figure 3.**
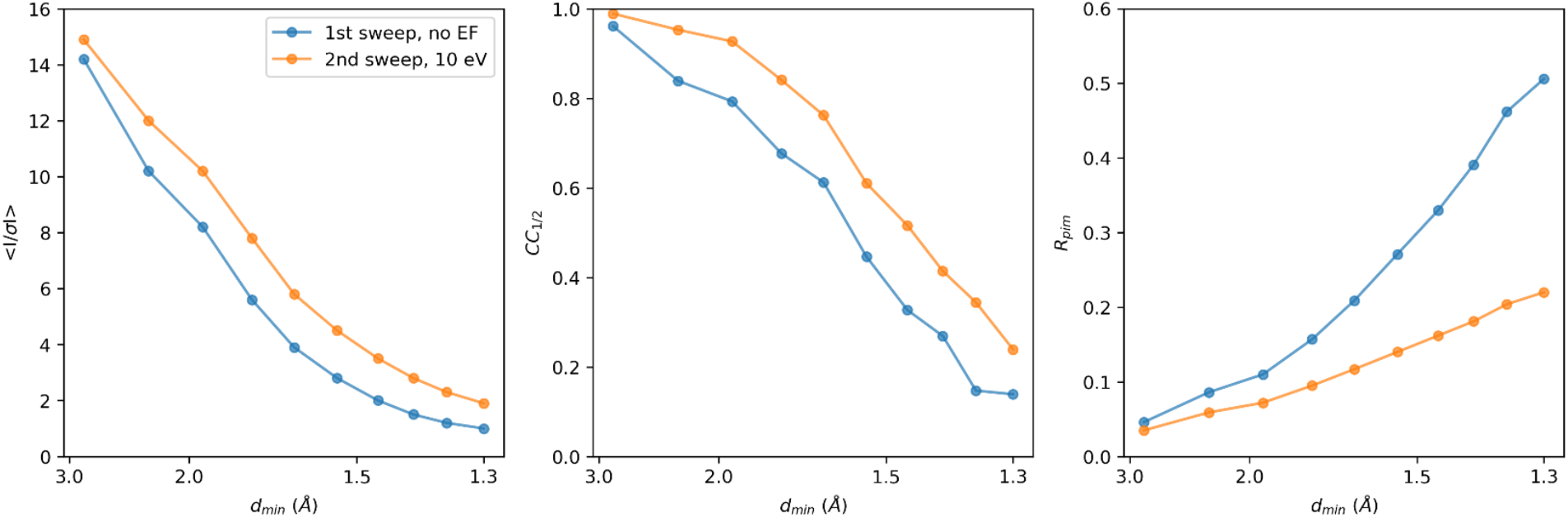
Merging statistics for unfiltered and filtered MicroED data. The crystallographic quality indicators mean *I/σI, CC*_1/2_, and *R*_pim_ are plotted as function of the resolution. For a side-by-side comparison, both data were merged and truncated at 1.3 Å resolution.

Our results highlight the advantages of energy filtering in MicroED for macromolecular crystallography. By directly comparing filtered and unfiltered data recorded from two sweeps of the same proteinase K lamellae, we demonstrate that energy filtering recovers high-resolution information, even after the crystal has absorbed double the amount of radiation dose. These improvements translate into better intensity statistics and greater internal consistency. Importantly, our results further suggest that high-resolution reflections, which would otherwise have been lost due to dose-sensitivity, could be partially recovered using energy filtering which reduces the noise. While model quality was similar in these experiments, previous comparisons between unfiltered and filtered data collected from separate but comparable crystals at the same fluence showed improve data quality with energy filtering (Clabbers *et al*., 2024). Together, these findings underscore the potential of energy filtering to overcome key limitations in electron crystallography, and indeed all cryo-EM modalities, offering a robust approach that reduces noise and partly mitigates information loss, yielding higher-resolution structures with improved reliability and precision.

## Methods

### Crystallization

Proteinase K microcrystals were are grown as previously described (Masuda *et al*., 2017; Clabbers *et al*., 2025a). Briefly, crystals were grown by mixing 40 mg/ml protein solution in 20 mM MES-NaOH pH 6.5 at a 1:1 ratio with a precipitant solution of 0.5 M NaNO_3_, 0.1 M CaCl_2_, 0.1 M MES-NaOH pH 6.5. The mixture was incubated at 4 °C, and microcrystals with dimensions of approximately 7-12 μm appeared within 24h.

### Sample preparation

A standard holey carbon electron microscopy grid (Quantifoil, Cu 200 mesh, R2/2) was glow discharged for 30 s at 15 mA on the negative setting. The sample was prepared using a Leica GP2 vitrification device set at 4 °C and 90% humidity, where 3 μl of crystal solution was deposited onto the grid, incubated for 10s, and any excess liquid was blotted away from the back side. The sample was then soaked on-grid for 30s with 3 μl cryoprotectant solution of 30% glycerol, 250 mM NaNO_3_, 50 mM CaCl_2_, 60 mM MES-NaOH pH 6.5. Any excess solution was blotted away using filter paper, and the grid was rapidly vitrified using liquid ethane.

### Focused ion beam milling

The grid was loaded onto a Helios Hydra 5 CX dual-beam plasma FIB/SEM (Thermo Fisher Scientific). Prior to milling, the grid was coated with a thin protective layer of platinum for 45s using the gas injection system. Microcrystals of proteinase K were machined using a 30 kV Argon plasma ion beam with a stepwise protocol as described previously (Martynowycz *et al*., 2023; Clabbers *et al*., 2024). Briefly, coarse milling steps were performed using a 2.0 nA current to a thickness of approximately 3 μm. Finer milling steps at 0.2 nA were used to thin the lamellae to 600 nm. Final polishing steps were performed at 60 pA down to an optimal thickness of 300 nm, equal to approximately one time the inelastic mean free at 300 kV (Martynowycz *et al*., 2021). In one session, 11 lamellae were prepared sequentially using this same protocol on the same grid. After milling, the grid was directly cryo-transferred to the transmission electron microscope (TEM) for data collection. During the transfer step, the grid was rotated by 90° relative to the milling direction such that the rotation axis on the microscope is perpendicular to the milling direction.

### Data collection

Diffraction data were collected on a Titan Krios G3i TEM (Thermo Fisher Scientific) equipped with a X-FEG operated at an acceleration voltage of 300 kV, a post-column Selectris energy filter, and a Falcon 4i direct electron detector. The microscope was aligned for low flux density conditions using the 50 μm C2 aperture, spot size 11, gun lens setting 8, and a parallel electron beam of 10 μm diameter. Under these conditions, the flux density was approximately 0.002 e^-^/Å^2^/s. The energy spread of the emitted electrons was characterized as *ΔE* = 0.834 eV +/-0.006 eV at full-width half max (FWHM). The zero-loss peak of the energy filter was aligned in defocused diffraction mode and centered within the SA aperture. Data were collected using the 150 μm selected area (SA) aperture, defining an area with ∼3.5 μm diameter at the sample plane. The detector distance was 1402 mm and calibrated prior to data collection using a standard evaporated aluminum grid (Ted Pella). For each lamella, MicroED data were collected using the continuous rotation method (CITE) over two consecutive passes. The first pass involved unfiltered data collection with the energy filter slit retracted, covering a 20.0° rotation range with an angular increment of 0.0476 °/s, an exposure time of 420 s, and a total fluence of 0.82 e-/Å2 (equal to an absorbed dose of ∼3.1 MGy). The second pass followed immediately after, using the same protocol but with the 10 eV energy filter slit inserted. The total absorbed dose after the second sweep was approximately 6.2 MGy. Equivalent dose values were calculated using the EMED subprogram of RADDOSE-3D (Dickerson *et al*., 2024). Data were recorded on a Falcon 4i direct electron detector in electron counting mode, operating at an internal frame rate of ∼320 Hz. The proactive dose protector was manually disabled. Raw data were written in electron event representation (EER) format with an effective readout speed of ∼308 frames per second, not counting gap frames.

### Data processing and refinement

Individual MicroED datasets in EER format were binned by two and converted to SMV format after applying post-counting gain corrections using the MicroED tools (available at https://cryoem.ucla.edu/downloads). Diffraction data were summed in batches of 308, such that each summed image represents a 1 s exposure. Individual MicroED datasets were processed using XDS (Kabsch, 2010). Data were integrated up to a cross-correlation between two random half sets that was still significant at the 0.1% level (Karplus & Diederichs, 2012). Individual datasets were merged using XSCALE (Kabsch, 2010). The merged data were truncated at a 1.3 Å resolution, enabling a direct comparison between intensity and model statistics. Data were merged using Aimless (Evans & Murshudov, 2013) and refined against the same proteinase K model (PDB ID 9DHO), which we reported previously at 1.09 Å resolution from energy-filtered MicroED data (Clabbers *et al*., 2025a). Both structures were refined using the same protocol in phenix.refine (Afonine *et al*., 2012) including electron scattering factors, automated optimization of the geometry and ADP weights, and individual anisotropic *B*-factors for all non-hydrogen atoms.

## Acknowledgements

This study was supported by the National Institutes of Health (P41GM136508), the Department of Defense (HDTRA1-21-1-0004), and the Howard Hughes Medical Institute. This study was supported by the National Institutes of Health P41GM136508. Portions of this research or manuscript completion were developed with funding from the Department of Defense MCDC-2202-002. Effort sponsored by the U.S. Government under Other Transaction number W15QKN-16-9-1002 between the MCDC, and the Government. The US Government is authorized to reproduce and distribute reprints for Governmental purposes, notwithstanding any copyright notation thereon. The views and conclusions contained herein are those of the authors and should not be interpreted as necessarily representing the official policies or endorsements, either expressed or implied, of the U.S. Government. The PAH shall flow down these requirements to its sub awardees, at all tiers.

## Data availability

Coordinates and structure factors have been deposited to the PDB.

---

**Figure S1.**
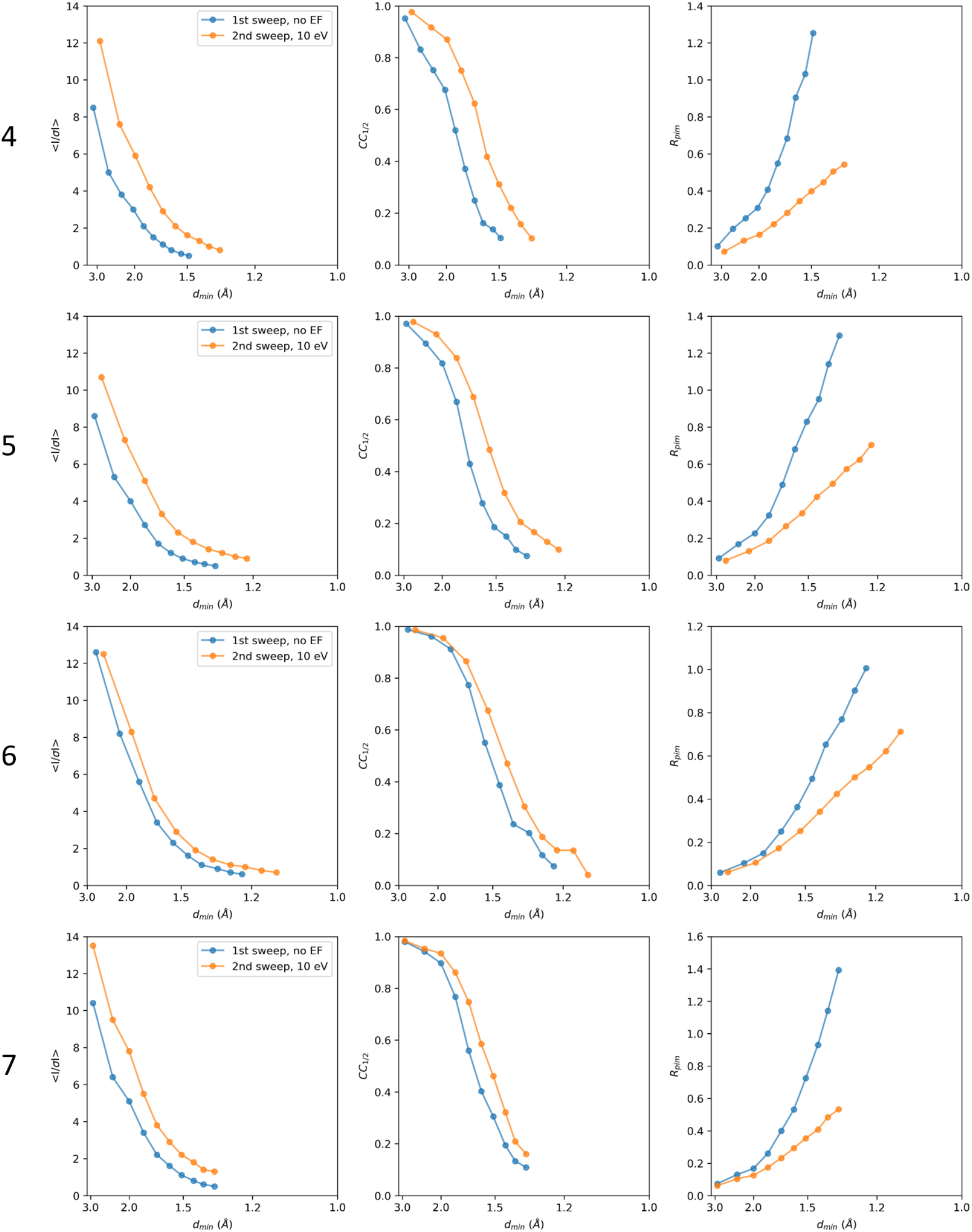
Intensity statistics for four MicroED data collection series. For each series, two datasets were

**Figure S2.**
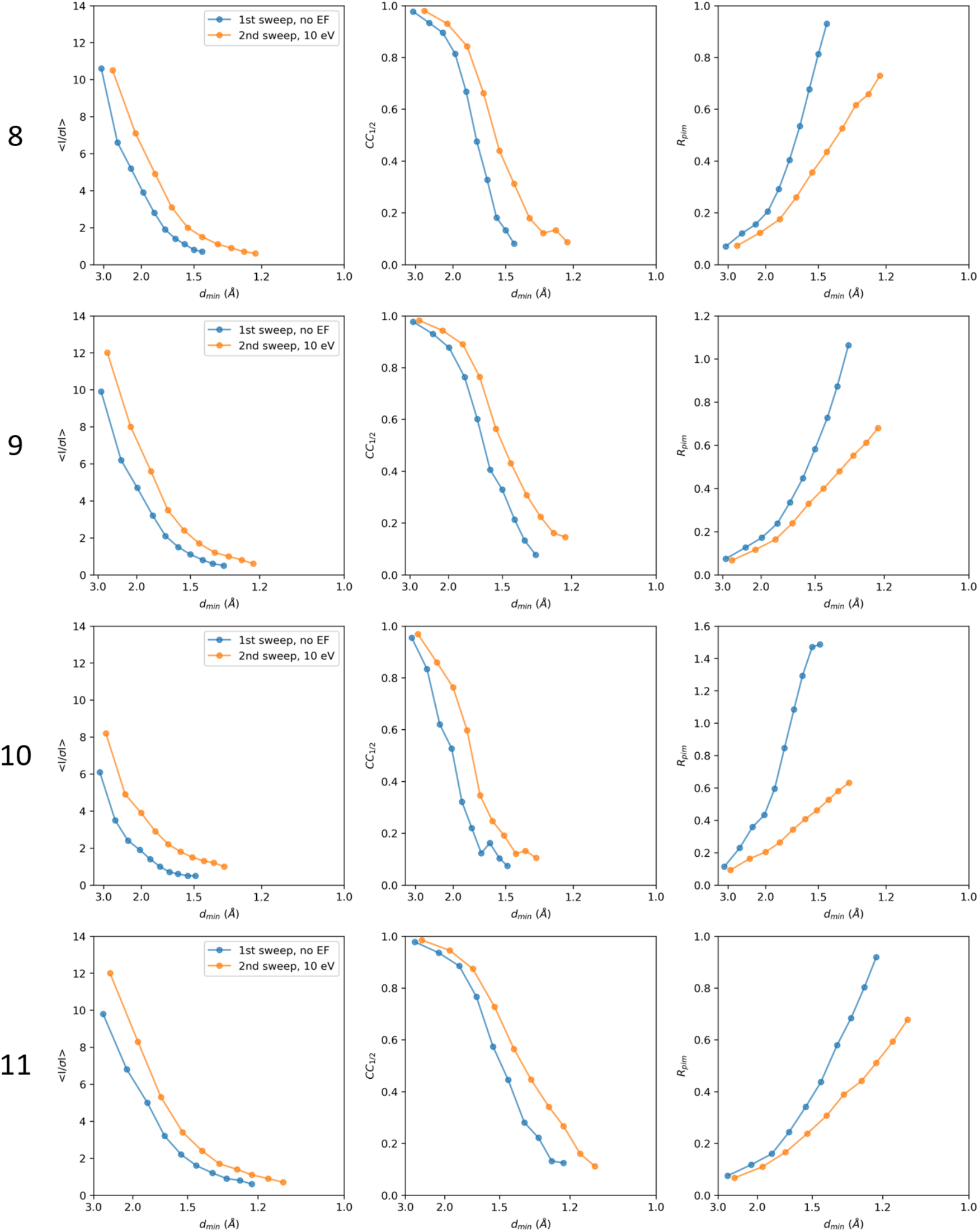
Intensity statistics for four MicroED data collection series. For each series, two datasets were

## Notes

### Competing Interest Statement

The authors have declared no competing interest.

## References

Afonine, P. V., Grosse-Kunstleve, R. W., Echols, N., Headd, J. J., Moriarty, N. W., Mustyakimov, M., Terwilliger, T. C., Urzhumtsev, A., Zwart, P. H. & Adams, P. D. (2012). Acta Crystallogr. D Biol. Crystallogr. 68, 352–367.

Clabbers, M.T.B., Hattne, J. Martynowycz, M.W. & Gonen, T. (2025). Nat. Commun. 16, 2247.

Clabbers, M.T.B., Hattne, J. Martynowycz, M.W. & Gonen, T. (2025). Preprint at bioRxiv, 10.1101/2025.02.24.639939.

Dickerson, J. L., McCubbin, P. T. N., Brooks-Bartlett, J. C. & Garman, E. F. (2024) Protein Sci. 33, e5005.

Evans, P. R. & Murshudov, G. N. (2013). Acta Crystallogr. D Biol. Crystallogr. 69, 1204–1214.

Garman, E.F. (2010). Acta Crystallogr. D Biol. Crystallogr. 66, 339–351.

Gemmi, M. & Oleynikov, P. (2013). Z. Für Krist. - Cryst. Mater. 228, 51–58.

Gonen, T., Sliz, P., Kistler, J., Cheng, Y. & Walz, T. (2004). Nature. 429, 193–197.

Gonen, T., Cheng, Y., Sliz, P., Hiroaki, Y., Fujiyoshi, Y., Harrison, S. C. & Walz, T. (2005). Nature. 438, 633–638.

Hattne, J., Shi, D., Glynn, C., Zee, C.-T., Gallagher-Jones, M., Martynowycz, M. W., Rodriguez, J. A. & Gonen, T. (2018). Structure 26, 759-766.e4.

Henderson, R. (1995). Q. Rev. Biophys. 28, 171–193.

Kabsch, W. (2010). Acta Crystallogr. D Biol. Crystallogr. 66, 125–132.

Karplus, P. A. & Diederichs, K. (2012). Science 336, 1030–1033.

Latychevskaia, T. & Abrahams, J. P. (2019). Acta Crystallogr. Sect. B Struct. Sci. Cryst. Eng. Mater. 75, 523–531.

Martynowycz, M. W., Clabbers, M. T. B., Hattne, J. & Gonen, T. (2022). Nat. Methods 19, 724–729.

Martynowycz, M. W., Clabbers, M. T. B., Unge, J., Hattne, J. & Gonen, T. (2021). Proc. Natl. Acad. Sci. 118, e2108884118.

Martynowycz, M. W., Shiriaeva, A., Clabbers, M. T. B., Nicolas, W. J., Weaver, S. J., Hattne, J. & Gonen, T. (2023). Nat. Commun. 14, 1086.

Masuda, T., Suzuki, M., Inoue, S., Song, C., Nakane, T., Nango, E., Tanaka, R., Tono, K., Joti, Y., Kameshima, T., Hatsui, T., Yabashi, M., Mikami, B., Nureki, O., Numata, K., Iwata, S. & Sugahara, M. (2017). Sci. Rep. 7, 45604.

Yang, T., Xu, H. & Zou, X. (2022). J. Appl. Crystallogr. 55, 1583–1591.

Yonekura, K., Ishikawa, T. & Maki-Yonekura, S. (2019). J. Struct. Biol. 206, 243–253.

Yonekura, K., Kato, K., Ogasawara, M., Tomita, M. & Toyoshima, C. (2015). Proc. Natl. Acad. Sci. 112, 3368–3373.

Yonekura, K., Maki-Yonekura, S. & Namba, K. (2002). Biophys. J. 82, 2784–2797.

